# Emergence of Lyme disease on treeless islands in Scotland, UK

**DOI:** 10.1101/2020.08.31.263319

**Authors:** Caroline Millins, Walter Leo, Isabell MacInnes, Johanne Ferguson, Graham Charlesworth, Donald Nayar, Reece Davison, Jonathan Yardley, Elizabeth Kilbride, Selene Huntley, Lucy Gilbert, Mafalda Viana, Paul Johnson, Roman Biek

## Abstract

Lyme disease (LD) is typically associated with forested habitats but has recently emerged on treeless islands in the Western Isles of Scotland. This has created a need to understand the environmental and human components of LD risk in open habitats. This study quantified both elements of LD risk and compared these between treeless islands with high and low LD incidence. We found high LD incidence was linked to higher prevalence in ticks (6.4% vs 0.4%) and increased human tick bite exposure. Most reported tick bites (72.7%) were within 1km of the home address and commonly in gardens. Residents on islands with high LD incidence reported increasing problems with ticks and suggested changing deer distribution as a potential driver. This study highlights the benefits to public health of an integrated approach to understand the factors contributing to LD emergence and a need to evaluate LD ecology in treeless habitats more broadly.

## Introduction

To optimize public health responses to vector-borne disease emergence, knowledge of the factors affecting the density of infected vectors (environmental hazard) in different habitats, interactions of people with this underlying hazard and how this relates to disease incidence are essential. Lyme disease (LD) is considered the most important emerging vector-borne zoonotic disease in both Europe and North America (1,2) and higher environmental hazard, LD cases and increased LD incidence are linked to wooded habitats (3–5). However, the recent emergence of LD on treeless islands in Scotland (6) has challenged our current understanding of the associations between habitat and LD incidence for this vector-borne disease. Within the UK, LD is an emerging zoonosis with the highest incidence reported from the Highland region of Scotland (7,8). Some islands in the Highland region with no woodland cover have a LD incidence twenty times the national average (119 cases/100,000 vs 3.2 cases/100,000) (6). A higher incidence on these islands has been recorded for a decade, while nearby and ecologically similar islands have a much lower incidence of 8.3 cases/100,000 (6). To understand, predict and mitigate LD emergence in treeless habitats, knowledge of the factors affecting the environmental hazard, how people interact with this hazard and possible drivers of emergence are urgently needed.

Though comparatively less is known about the environmental hazard from ticks in treeless habitats, available evidence suggests that in general, the hazard is lower than in wooded areas. Woodlands are widely regarded as the optimum habitats for the Ixodid tick vector due to the suitably humid microclimate which improves off-host tick survival and availability of hosts for blood meals (9,10). Studies which sample ticks from woodland and nearby grassland commonly find lower tick density in grassland, leading to suggestions that grassland may act as a ‘sink’ for tick populations maintained by more suitable conditions in woodlands (11–13). In general, studies of tick populations in treeless habitats have found that the density of *Ixodes ricinus*, the main tick vector of LD, tends to be much lower compared to woodlands (14). For example, surveys of open habitats in northern Spain found no questing *I. ricinus* (15), and within the UK most studies have found a relatively low density in meadows (16), open hillside (17,18) and heather moorland (19–21).

In the absence of longitudinal environmental and tick abundance data in treeless areas impacted by LD emergence, alternative approaches are needed to assess if changes in tick population abundance and distribution have occurred over time. Tick populations in treeless habitats will be subject to many of the same environmental drivers as those in forested areas, including changes in climate, land management and host density, particularly deer populations (22–25). Surveys of local communities can provide information on whether the tick hazard is perceived to have changed over time, and suggest possible environmental factors associated with this change which can focus future research (26).

Environmental hazard is linked to LD incidence by how people interact with the environment and become exposed to infected tick bites (27). People’s activities, knowledge and attitudes towards the risk of tick bites, and behaviours taken to reduce tick bites before entering tick habitats will influence tick bite exposure (27,28) Knowledge of where people become exposed to tick bites, risk factors associated with tick bite exposure and the environmental hazard in different habitats can then be used to direct preventative public health interventions (29).

To assess the ecological and human dimensions contributing to high LD incidence in treeless habitats and identify possible drivers of emergence we assessed i) factors influencing tick density and prevalence of *B. burgdorferi* s.l. (components of the environmental hazard) ii) community observations of whether problems with ticks have changed over time and iii) identified geographical locations where people are exposed to tick bites and demographic and behavioural factors affecting tick bite exposure, using treeless islands with high and low LD incidence in the Western Isles in Scotland, UK as our study system.

## Methods

### Site Selection

To compare the environmental hazard between islands experiencing high LD incidence (North Uist, South Uist and Benbecula; 26 sites), and low incidence (Harris/Lewis and Barra; 16 sites), we selected sites within two dominant habitat types, improved grassland and heather moorland, using a spatially stratified sampling design and the random selection tool in QGIS (QGIS Development Team, 2018) (Figure 1).

**Figure 1.**
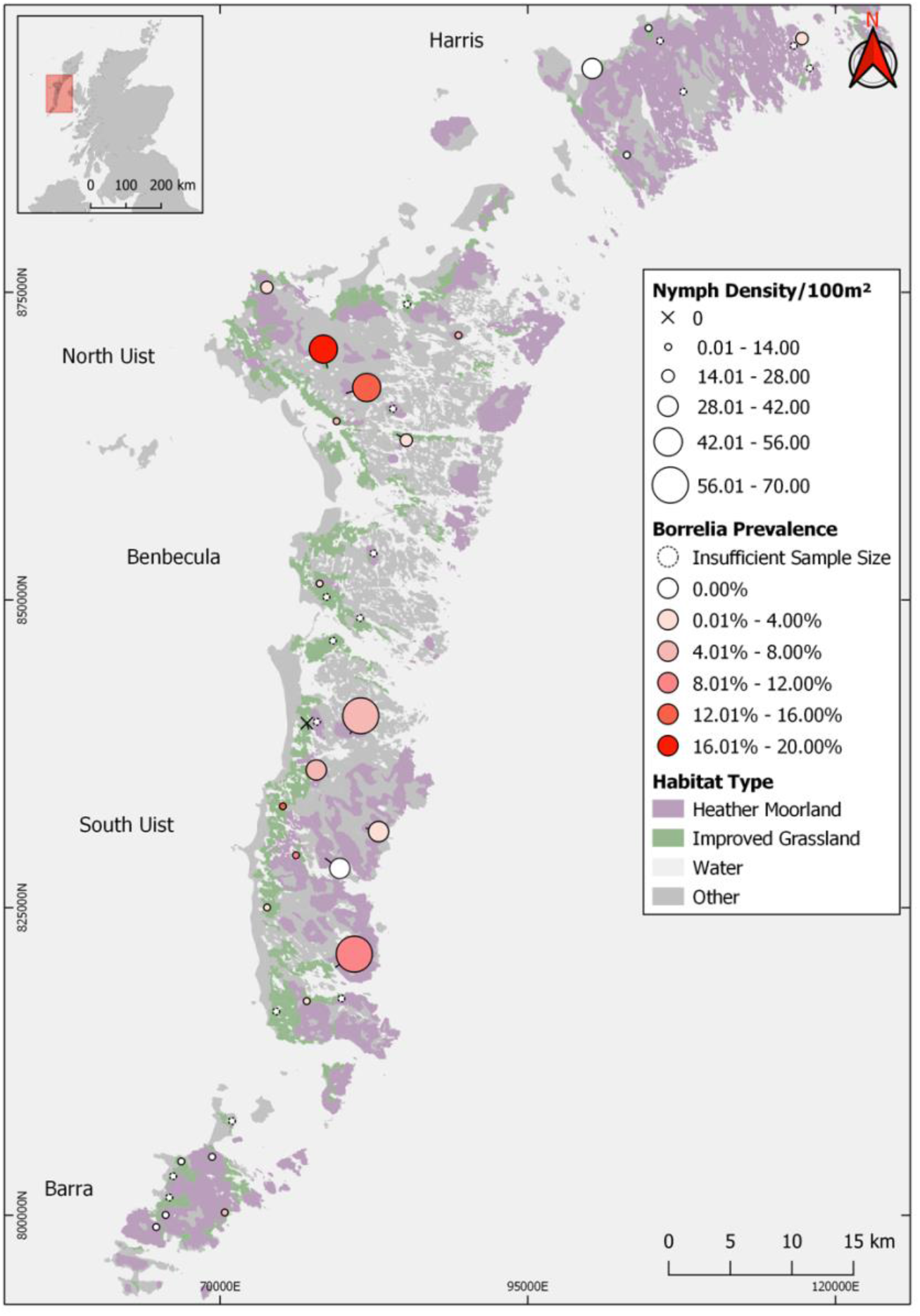
Map to show study sites in improved grassland and heather moorland in high (North and South Uist, Benbecula) and low Lyme disease incidence islands (Harris, Barra), Western Isles, UK sampled in 2018-2019. Circle sizes are in proportion to the questing tick density and *Borrelia burgdorferi* sensu lato prevalence is represented by the graded colour of the circles. Prevalence was not estimated at sites where < 50 ticks were collected, sites at which no ticks were detected are shown as a ‘x’.

**Figure 2.**
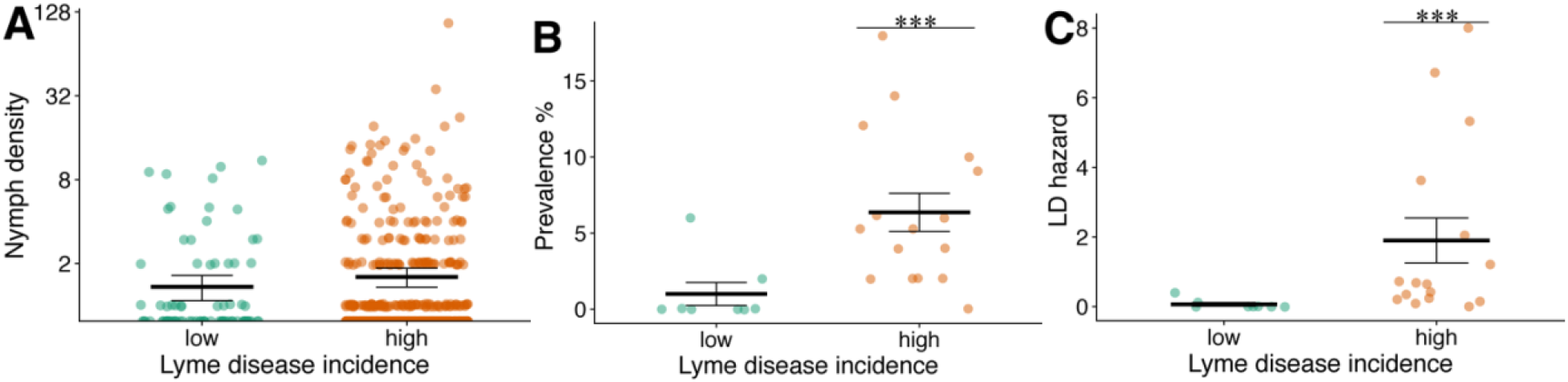
Comparison of mean tick density per 10 m^2^ blanket drag (A), prevalence of *B. burgdorferi* s.l. per site (B), LD hazard (infected nymphs/100 m^2^) among high and low Lyme disease incidence island grassland and moorland sites shown in Figure 1, Western Isles, UK. Horizontal bars represent means with standard errors. High and low LD incidence sites differed significantly with respect to prevalence and LD hazard, but not nymph density (Table 1).

**Figure 3.**
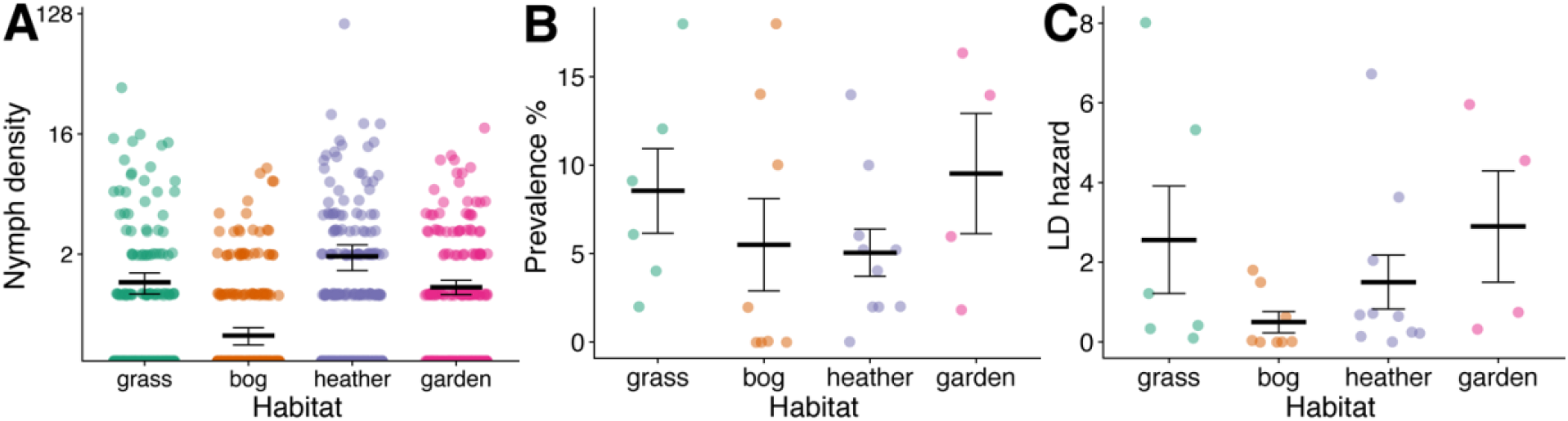
Comparison of mean tick density per 10 m^2^ blanket drag (A), prevalence of *B. burgdorferi* s.l. per site (B) and LD hazard (infected nymphs/100 m^2^) (C) among different habitat types in high Lyme disease incidence islands. Improved grassland (grass), Bog and peatland (bog), Heather moorland (heather) and garden sites, Western Isles, UK (Machair sites not shown due to very low mean tick density = 0.025 nymphs/10 m^2^ (+/− SE 0.01-0.04). Standard error bars are shown on plots. No statistically significant differences in tick density, prevalence and LD hazard among these four different habitat types were found (Appendix Table 2).

On islands with high LD incidence, sites were selected from three additional habitats, machair grassland (8 sites) and bog/peatland (13 sites) using the same stratified sampling approach and from domestic gardens (12 sites) randomly selected from each sector (Appendix Figure 1). Machair is a sandy grassland present along ocean coastline often used for grazing or cultivation (30). Sites were sampled during the peak questing period for *I. ricinus* from April 19th, 2018 – June 5th, 2018. Additional sites from low LD incidence areas were sampled from 17^th^ May, 2019 – 22^nd^ June, 2019 to strengthen the comparison of tick infection prevalence.

### Tick Collection

To estimate questing *I. ricinus* density, twenty randomized transects 10 m in length were sampled at each site, with transects 30-50 m apart, or 20-30 m apart in gardens. At the start of each transect, vegetation height and density and temperature and humidity were measured (31). A 1 m^2^ white woollen blanket was dragged across the surface of the vegetation for 10m and questing nymphs were collected on the blanket, counted and placed in 100% ethanol. Continuous blanket dragging was carried out for up to two person hours to increase the sample size of ticks collected at each site.

### Screening of *I. ricinus* ticks for *B. burgdorferi* sensu lato and genospecies identification

*I. ricinus* nymphs were extracted individually, up to a target sample size of approximately 50 nymphs at each site, using an ammonia hydroxide extraction technique (32). A target sample size of 50 nymphs was selected as the minimum sample size to detect an infected tick based on the expected prevalence from a pilot study of 6.6%. A nested PCR which targeted the flagellin gene was used to test for *B. burgdorferi* s.l. with sequencing of the 390 base-pair product (33) to identify the genospecies.

### Geographical locations of human tick bite exposure, factors associated with tick bite risk and perceptions of tick problems over time

A questionnaire was used to collect data about tick bite exposure among residents. The survey was designed to test if there was a difference in tick bite exposure between high and low LD incidence areas, identify habitat types where tick bites occurred, the distance of tick bites relative to the home address, and social and behavioural factors associated with exposure to tick-bites. Residents were asked if problems with ticks had changed over time. The survey was approved by the University of Glasgow College of Medical, Veterinary and Life Sciences Ethics Committee (reference number 200170121) and available 18 April 2018 – 31st October 2018.

### Statistical Analysis

All statistical analyses and model selection were carried out in R version 4.0.0 (R Development Core Team, Vienna, Austria) using the *lme4* package for generalized linear mixed models (GLMMs)(34). For all models, correlations between explanatory variables were tested for using the variance inflation function (VIF) in the *car* package in R (35). Overdispersion was assessed by calculating the ratio of the sum of squared Pearson residuals to the number of residual degrees of freedom. A backwards model selection process based on minimizing AIC was carried out starting from the maximum global model, variables were retained which improved model fit (delta AIC > 2) (36). The change in AIC (delta AIC) caused by removing each explanatory variable from the final model in turn was calculated (37). The *lsmeans* package in R was used to make post hoc pairwise comparisons within categorical explanatory variables (38).

To test for an association between tick density and LD incidence (high/low) we modelled nymph abundance (count of nymphs per 10 m drag), using a GLMM with a log link and Poisson distributed errors, as a function of the categorical variables LD incidence (high, low), habitat type and wind (on Beaufort wind force scale), and the continuous variables vegetation density, temperature and humidity, with random effects of site and observation (39). Within islands of high LD incidence, where we sampled additional habitat types, we used a separate model to test the effect of habitat type and island on tick abundance with the same covariates.

To test for an association between tick infection prevalence with *B. burgdorferi* s.l. and LD incidence (high or low) we modelled nymph prevalence at sites which had a minimum of approximately 50 nymphs collected. We modelled the proportion of infected nymphs using a GLMM with a logit link and binomially distributed errors as a function of LD incidence (high/low), habitat type, mean nymph density and with a random effect of site. Within islands of high LD incidence where additional habitat types were sampled, we used a separate model to test for the effect of habitat and island on tick prevalence. Machair grassland was not included in analyses due to low tick densities and sample sizes at all sites.

To test for an association between the environmental LD hazard (density of infected ticks) and LD incidence (high/low), we modelled the number of infected ticks using a GLMM with a log link and Poisson distributed errors as a function of LD incidence (high or low) and habitat, with an offset of the log of the estimated area surveyed to collect the total number of ticks which were tested (which converted the predicted outcome to the number of infected ticks per m^2^), and with a random effect of site. Separate GLMM models were used to test for the effect of habitat and island on LD hazard within high LD incidence islands.

Responses from a community survey were analysed to determine if problems with ticks and tick-borne diseases had changed over time. A total of 522 surveys were received from adult residents of the Western Isles and used for the analysis. We hypothesized that a higher proportion of respondents in areas of high LD incidence areas would report in their comments that tick numbers and associated problems were increasing. Text responses were categorized as ‘increased’ or ‘not increased’ and used in a GLM with a logit link and binomially distributed errors with LD incidence (high, low) as an explanatory variable. Themes present in free text responses were compared between areas with high and low LD incidence to assess factors associated by the community with emergence of problems associated with ticks. A corpus linguistic approach was used to extract common keywords and associated clusters of words to explore the meaning (Appendix Methods, 40).

To test for differences in human tick bite exposure between areas with high and low LD incidence, community survey responses (n = 522) were used. Tick bite exposure classified as high risk (≥ 5 tick bites per year) or low risk (< 5 tick bites per year) was modelled with a GLM, a logit link and binomially distributed errors as a function of the following variables: LD Incidence (high, low), age, gender, occupation risk, frequency of outdoor activity and pet ownership (all categorical variables). Variables were tested by univariable analysis to maximise sample size for each and to test for interactions with LD incidence (high, low) prior to inclusion in a multivariable model. As awareness, attitudes and prevention of tick bites could influence reported tick-bite exposure, we tested for associations between tick bite exposure and these variables in a separate model and included an interaction of each variable with LD Incidence (high, low).

It was hypothesized that tick presence in a home could result from ticks being transported on people’s clothing or on pets and this could be associated with LD Incidence (high/low). To test this, we used survey responses to determine whether any tick (live unfed, engorged or dead) had ever been detected inside the home and modelled this with a GLM, a logit link and binomially distributed errors as a function of; LD Incidence, level of outdoor activity and pet ownership.

## Results

### Tick density

Mean tick density per 10 m^2^ at low LD incidence improved grassland and heather moorland sites sampled in 2018 was 1.36 nymphs (+/−SE =1.08-1.64) compared to 1.60 nymphs (+/−SE =1.35-1.85) from the same habitats sampled from high LD incidence sites (Figure 1, Appendix Table 1). Tick density did not vary with LD incidence (high, low), which was dropped from the final model (Table 1). Within sites located on high LD incidence islands (Appendix Figure 1, Appendix Table 1), fixed effects selected in the best fit model included habitat type (delta AIC = 16.06, Appendix Table 2). Significantly fewer nymphs were found in machair grassland compared to all other habitat types (Tukey’s post hoc comparison p<0.01); there were no significant differences in nymph density between other habitat types.

**Table 1.** Results of best fit generalized linear mixed model to explain variation in questing nymph density, nymph prevalence and Lyme disease (LD) hazard (density of infected nymphs) between high and low LD incidence sites.

| Model | Variable | Estimate | Standard<br>error | p-value | delta AIC* |
| --- | --- | --- | --- | --- | --- |
| Tick density | (Intercept) | -4.02 | 0.93 | <0.001 | NA |
|  | Temperature<br>(°C/10) | 2.11 | 0.64 | <0.001 | 9.44 |
| Tick prevalence | (Intercept) | -5.19 | 0.59 | <0.001 | NA |
|  | LD Incidence |  |  |  |  |
|  | Low | Reference |  |  |  |
|  | High | 2.33 | 0.61 | <0.001 | 13.5 |
| LD hazard | (Intercept) | -8.22 | 0.96 | <0.001 | NA |
|  | LD Incidence |  |  |  |  |
|  | Low | Reference |  |  |  |
|  | High | 3.57 | 0.91 | <0.001 | 14.34 |

\*delta AIC is the change in AIC from removing each variable from the best fit model.

### *B. burgdorferi* s.l. prevalence

The prevalence of *B. burgdorferi* s.l. in high LD incidence sites was significantly higher (6.43% (+/−SE 5.61% – 7.26%, n=57/886) compared to low LD incidence sites (0.66%: +/− SE 0.33%-0.98%, n=4/609 nymphs) (Table 1). In high LD incidence sites, 98.25% of infected nymphs carried *B. afzelii* (56/57 infected nymphs), and 1.75% (1/57 infected nymphs) *B. garinii*. In low LD incidence sites, 75% of infected nymphs carried *B. garinii* (3/4 infected nymphs) and 25% carried *B. valaisiana* (1/4 infected nymphs).

Within high LD incidence sites, prevalence did not differ by island or habitat type; an intercept model was the final model in both cases (Appendix Table 2).

### LD hazard

A significantly higher mean LD hazard (1.90 nymphs per 100 m^2^ (+/− SE 1.25-2.55) was found in high LD incidence sites compared to low LD incidence sites (0.07 +/−SE 0.02-0.12). Within high LD incidence sites, hazard did not differ by island or habitat type, an intercept model was the final model in both cases (Appendix Table 2).

### Geographical locations of tick bite risk

Most participants (64.4%, n=333/517) provided information on the habitat type of their last tick bite and island of residence (Appendix Results). Of respondents providing tick bite information, 51.7% (n=172/333) provided a spatial location which enabled the distance of the tick bite from their residence to be calculated. This revealed that 72.7% (125/172) of tick bites occurred within 1 km of the home address, and 47.1% (81/172) were at the home address (Appendix Figure 2).

### Factors associated with tick bite exposure risk

The effects of LD incidence (high, low) and other factors known to affect tick bite exposure risk were examined individually (Appendix Table 3) and in a multivariable general linear model. The most significant factor in a multivariable model explaining tick bite exposure risk was LD incidence, with residents from areas of higher LD incidence reporting higher rates of tick bite exposure compared to lower LD incidence areas (OR = 2.20, 95% CI 1.37 – 3.60: Table 2). No difference in awareness, attitudes and prevention was found between high and low incidence LD islands.

**Table 2.** Results of the best fit general linear model to test for factors affecting risk of tick bite exposure in residents of the Western Isles, UK, classified as high (≥5 tick bites per year) or low (< 5 tick bites per year).

| **Variable** | **Estimate** | **Standard**<br>**error** | **p value** | **Odds ratio**<br>**(95% CI)** | **delta AIC\*** |
| --- | --- | --- | --- | --- | --- |
| **Intercept** | -2.75 | 0.52 | <0.001 | NA | NA |
| **LD† incidence** |  |  |  |  |  |
| **Low** | (reference) |  |  | 1.0 (reference) |  |
| **High** | 0.79 | 0.25 | 0.001 | 2.20 (1.37-3.60) | 8.64 |
| **Age** |  |  |  |  |  |
| **18-30** | (reference) |  |  | 1.0 (reference) |  |
| **30-60** | 0.67 | 0.44 | 0.132 | 1.95 (0.86-5.01) |  |
| **>60** | 1.28 | 0.48 | 0.008 | 3.58 (1.46-9.82) | 4.76 |
| **Outdoor activity** |  |  |  |  |  |
| **Less than most days** | (reference) |  |  | 1.0 (reference) |  |
| **Most days** | 0.63 | 0.28 | 0.025 | 1.87 (1.10-3.29) | 3.4 |

†Lyme disease (LD).
\* delta AIC is the change in AIC from removing each variable from the best fit model

### Factors associated with finding a tick within the home

The chances of finding a tick within the home were not affected by LD incidence but increased with pet ownership (OR 4.07, 95% C.I. 2.61-6.41, delta AIC = 36.94), and increased slightly with ‘outdoor activity most days’ (OR 1.67, 1.05-2.64, delta AIC = 2.77; Appendix Table 4).

### Changes in tick numbers and problems over time

Overall, the majority of respondents (50.6%, 210/415) described problems with ticks increasing over time. Residents from high LD incidence islands were significantly more likely to say that tick numbers and associated problems had increased overtime (OR 4.5, 95% CI 2.1 – 10.0), p < 0.001, delta AIC 13.5 (Appendix Table 5). Linguistic analysis of free text comments revealed differences in themes between high and low LD incidence islands. More ticks were reported to be present across all islands, with responses from high LD incidence islands only showing stronger emphasis for example, ‘definitely increased’ or ‘significantly increased’. Comments supported more deer in proximity to homes only on high LD incidence islands (Appendix Table 6).

## Discussion

Here we detail the first investigation of LD emergence in treeless habitats in Europe and show that both environmental hazard and human tick bite exposure contribute to higher LD incidence in these settings. In contrast to previous European studies, we found that the hazard amongst treeless habitats can be comparable to forested sites which are traditionally associated with higher LD risk (31,41).

Differences in the environmental hazard between high and low LD incidence locations was due to a significant difference in the prevalence of *B. burgdorferi* s.l. within questing ticks. Almost all infected ticks on high incidence islands carried *B. afzelii* a genospecies associated with mammalian transmission hosts (42) while no *B. afzelii* infection was detected in ticks collected from low incidence islands, where the prevalence was extremely low (<1%). Given the similarity in habitats and climate, we hypothesize that the presence/absence of this genospecies could either be driven by differences in the host community or that introduction of *B. afzelii* from the mainland has been limited to certain islands.

Within islands with a high incidence of LD, improved grassland, heather moorland, bog and peatland habitats and domestic gardens all had similar tick abundance and prevalence of *B. burgdorferi* s.l. as mainland forested sites in Scotland (31,41). Our results suggest that the microclimatic conditions in these open habitats, possibly driven by the milder oceanic climate on the Western Isles, can be as conducive to tick survival as in woodlands. Tick abundance was positively associated with vegetation density, which in combination with relatively high rainfall and humidity in this location, might contribute to a more favourable microclimate and improved off-host tick survival. In contrast we found significantly lower tick abundance within machair grassland, likely due to a combination of short vegetation height, lack of a vegetation mat and agricultural rotations and ploughing reducing off-host tick survival (43,44).

In common with previous studies, we found that in the absence of longitudinal data on vector populations and linked ecological drivers, community surveys can be valuable as an indicator (26). Ticks were significantly more likely to be reported as an increasing problem by residents living on islands with a high LD incidence, consistent with this being an emerging issue. Increased deer populations and deer presence near to homes, were commonly mentioned by affected communities as possible causes for increasing numbers of ticks. As an established driver of tick populations and distribution (22,45,46) and associated with LD emergence in other areas of Europe (47), the association between deer habitat use and tick density in the Western Isles should be investigated as part of future research.

In addition to a higher environmental hazard on high LD incidence islands, we found that residents on high LD incidence islands were more likely to report a higher exposure to tick bites. Tick bite exposure increased with age and amounts of outdoor activity but these factors and knowledge, attitudes and prevention of tick bites did not contribute to differences in tick-bite exposure between high and low LD incidence islands. Our work highlights the benefits of combining ecological surveys with data on human tick bite exposure. While, no significant differences in tick abundance on grassland and moorland sites were found between high and low LD incidence islands, questionnaire responses indicated that tick density in these habitats may not be the most relevant for human tick bite exposure in our study area. Most reported tick bites occurred close to the home address, and frequently in gardens. We found a similar environmental hazard in gardens to surrounding habitats on high LD incidence islands indicating that spill-over of infected ticks into peridomestic environments is common. Further research is required to test if peridomestic tick exposure may be driving differences in tick bite exposure between high and low LD incidence islands. Frequent tick bites within gardens and exposure of residents to tick bites within homes is of relevance to medical practitioners, as it suggests that all members of a household could be at risk from tick bites and develop LD. Our research suggests that targeted environmental and educational public health interventions focussed around people’s homes could reduce tick bite exposure and potentially cases of LD.

This study has shown that treeless habitats can support similar tick densities and infection risk as forested areas and can be associated with LD emergence in humans. The results from our study system suggest potential for LD to emerge in open habitats with a suitable microclimate for off-host tick survival and host availability for blood meals elsewhere in Europe. Integrating these results with human tick bite exposure revealed that the majority of tick bites occur close to homes, and that spillover of ticks and tick-borne pathogens into gardens and homes is an emerging problem which residents link to increased deer populations and their changing distribution. Further research to understand the effect of ecological drivers of tick populations in these regions, together with information on human use of these environments, will be necessary to achieve more accurate prediction of areas of risk and suggest ways to prevent and mitigate this risk.

## Acknowledgments

MV was funded by the European Research Council (ERC) under the European Union’s Horizon 2020 research and innovation programme (grant agreement No. 852957). JY was supported by a CASE studentship funded by the Natural Environment Research Council (NERC).

## Disclaimers

### Author Bio

Dr Caroline Millins is a Tenure Track Research Fellow in the Department of Livestock and One Health at the University of Liverpool. Her primary research interests include One Health approaches to the study of zoonotic pathogens, vector-borne pathogen ecology and wildlife health.

## References

1. Dennis D, Hayes E. Epidemiology of *Lyme Borreliosis*. In: Gray J, Kahl O, Lane R, Stanek G, editors. Lyme Borreliosis, Biology, Epidemiology and Control. New York: CABI; 2002. p. 251–80.

2. Rizzoli A, Hauffe HC, Carpi G, Vourc’h GI, Neteler M, Rosà R. Lyme borreliosis in Europe. Eurosurveillance [Online]. 2011;16(27):19906.

3. James MC. The ecology, genetic diversity and epidemiology of Lyme borreliosis in Scotland. PhD thesis. [Aberdeen]: University of Aberdeen; 2010.

4. Eisen RJ, Lane RS, Fritz CL, Eisen L. Spatial patterns of lyme disease risk in California based on disease incidence data and modeling of vector-tick exposure. Am J Trop Med Hyg. 2006;75(4):669–76.

5. Dister SW, Fish D, Bros SM, Frank DH, Wood AL. Landscape characterization of peridomestic risk for Lyme disease using satellite imagery. Am J Trop Med Hyg. 1997;57(6):687–92.

6. NHS-Western Isles. The ‘Tick’-ing time bomb. The incidence of Lyme disease in the Outer Hebrides (2010-2017) [Internet]. 5 Nations Health Protection Conference. 2018 [cited 2020 Aug 13]. Available from: https://www.wihb.scot.nhs.uk/wp-content/uploads/2020/08/A0-Template-The-ticking-time-bomb.-Incidence-of-Lyme-disease-in-the-Western-Isles-2010-2017.pdf

7. Slack GS, Mavin S, Yirrell D, Ho-Yen D. Is Tayside becoming a Scottish hotspot for Lyme borreliosis? J R Coll Physicians Edinb. 2011;41:5–8.

8. Tulloch JSP, Decraene V, Christley RM, Radford AD, Warner JC, Vivancos R. Characteristics and patient pathways of Lyme disease patients: A retrospective analysis of hospital episode data in England and Wales (1998-2015). BMC Public Health. 2019;19(1):1–11.

9. Medlock JM, Shuttleworth H, Copley V, Hansford K, Leach S. Woodland biodiversity management as a tool for reducing human exposure to *Ixodes ricinus* ticks: A preliminary study in an English woodland. J Vector Ecol. 2012;37(2):307–15.

10. Randolph SE. Tick ecology: processes and patterns behind the epidemiological risk posed by ixodid ticks as vectors. Parasitology. 2004 Oct;129(7):S37–65.

11. Boyard C, Barnouin J, Gasqui P, Vourc’h G. Local environmental factors characterizing *Ixodes ricinus* nymph abundance in grazed permanent pastures for cattle. Parasitology. 2007;134(7):987–94.

12. Hoch T, Monnet Y, Agoulon A. Influence of host migration between woodland and pasture on the population dynamics of the tick *Ixodes ricinus:* A modelling approach. Ecol Modell. 2010;221(15):1798–806.

13. Gilbert L. How landscapes shape Lyme borreliosis risk. In: Braks M, van Wieren SE, Takken W, Sprong H, editors. Ecology and Control of Vector Borne Diseases. Volume 4. Wageningen Academic Publishers (WAP).; 2016. p. 161–71.

14. Gray JS, Kahl O, Robertson JN, Daniel M, Estrada-Peña A, Gettinby G, et al. Lyme borreliosis habitat assessment. Zentralblatt fur Bakteriol. 1998;287(3):211–28.

15. Estrada-Pena A. Distribution, abundance, and habitat preferences of *Ixodes ricinus* (Acari: Ixodidae) in northern Spain. J Med Entomol. 2001;38:9.

16. Gray JS. The ecology of ticks transmitting Lyme borreliosis. Exp Appl Acarol. 1998;22:249–58.

17. Ogden NH, Nuttall PA, Randolph SE. Natural Lyme disease cycles maintained via sheep by co-feeding ticks. Parasitology. 1997;115:591–600.

18. Walker AR, Alberdi MP, Urquhart KA, Rose H. Risk factors in habitats of the tick *Ixodes ricinus* influencing human exposure to *Ehrlichia phagocytophila* bacteria. Med Vet Entomol. 2001;15(1):40–9.

19. Ruiz-Fons F, Gilbert L. The role of deer as vehicles to move ticks, *Ixodes ricinus*, between contrasting habitats. Int J Parasitol. 2010;40(9):1013–20.

20. Harrison A, Newey S, Gilbert L, Haydon DT, Thirgood S. Culling wildlife hosts to control disease: mountain hares, red grouse and louping ill virus. J Appl Ecol. 2010;47(4):926–30.

21. Gilbert L. Altitudinal patterns of tick and host abundance: a potential role for climate change in regulating tick-borne diseases? Oecologia (Berlin). 2010;162(1):217–25.

22. Medlock JM, Hansford KM, Bormane A, Derdakova M, Estrada-Peña A, George J-C, et al. Driving forces for changes in geographical distribution of *Ixodes ricinus* ticks in Europe. Parasit Vectors. 2013 Jan;6(1).

23. Werden L, Barker IK, Bowman J, Gonzales EK, Leighton PA, Lindsay LR, et al. Geography, Deer, and Host Biodiversity Shape the Pattern of Lyme Disease Emergence in the Thousand Islands Archipelago of Ontario, Canada. PLoS ONE [Online]. 2014 Jan;9(1):85640.

24. Ostfeld RS, Brunner JL. Climate change and Ixodes tick-borne diseases of humans. Philos Trans R Soc Lond B Biol Sci. 2015;370: 20140.

25. Simon J A., Marrotte RR, Desrosiers N, Fiset J, Gaitan J, Gonzalez A, et al. Climate change and habitat fragmentation drive the occurrence of *Borrelia burgdorferi*, the agent of Lyme disease, at the northeastern limit of its distribution. Evol Appl. 2014;7:750–64.

26. Kimaro EG, Toribio JALML, Mor SM. Climate change and cattle vector-borne diseases: Use of participatory epidemiology to investigate experiences in pastoral communities in Northern Tanzania. Prev Vet Med. 2017;147(August):79–89.

27. Bouchard C, Aenishaenslin C, Rees EE, Koffi JK, Pelcat Y, Ripoche M, et al. Integrated social-behavioral and ecological risk maps to prioritize local public health responses to lyme disease. Environ Health Perspect. 2018;126(4):1–13.

28. Fischhoff IR, Keesing F, Ostfeld RS. Risk Factors for Bites and Diseases Associated with Black-Legged Ticks: A Meta-Analysis. Am J Epidemiol. 2019;188(9):1742–50.

29. Finch C, Al-Damluji MS, Krause PJ, Niccolai L, Steeves T, O’Keefe CF, et al. Integrated assessment of behavioral and environmental risk factors for lyme disease infection on Block Island, Rhode Island. PLoS One. 2014;9(1).

30. Angus S. The Outer Hebrides: Moor and Machair. Winwick: White Horse Press; 2001.

31. Millins C, Gilbert L, Johnson P, James M, Kilbride E, Birtles R, et al. Heterogeneity in the abundance and distribution of *Ixodes ricinus* and *Borrelia burgdorferi* (sensu lato) in Scotland: implications for risk prediction. Parasites and Vectors. 2016;9(1).

32. Gern L, Douet V, López Z, Rais O, Cadenas FM. Diversity of Borrelia genospecies in *Ixodes ricinus* ticks in a Lyme borreliosis endemic area in Switzerland identified by using new probes for reverse line blotting. Ticks Tick Borne Dis. 2010 Mar;1(1):23–9.

33. Johnson BJB, Happ CM, Mayer LW, Piesman J. Detection of *Borrelia burgdorferi* in ticks by species-specific amplification of the flagellin gene. Am J Trop Med Hyg. 1992;47(6):730–41.

34. Bates D, Maechler M, Bolker B, Walker S. lme4: Linear mixed-effects, models using Eigen and S4. 2019.

35. Fox J, Weisberg S. An R Companion to Applied Regression. Third. Thousand Oaks CA: Sage; 2019.

36. Chambers JM. Chapter 4. Linear Models. In: Chambers JM, Hastie TJ, editors. Statistical models in S. Wadsworth & Brooks/Cole; 1992. p. 100–45.

37. Burnham KP, Anderson DR. Multimodel inference: Understanding AIC and BIC in model selection. Sociol Methods Res. 2004;33(2):261–304.

38. Lenth RV. Least-Squares Means: The {R} Package {lsmeans}. J Stat Softw. 2016;69(1):1–33.

39. Elston DA, Moss R, Boulinier T, Arrowsmith C, Lambin X. Analysis of aggregation, a worked example: numbers of ticks on red grouse chicks. Parasitology. 2001;122(Pt 5):563–9.

40. Huntley SJ, Mahlberg M, Wiegand V, van Gennip Y, Yang H, Dean RS, et al. Analysing the opinions of UK veterinarians on practice-based research using corpus linguistic and mathematical methods. Prev Vet Med. 2018;150(January 2017):60–9.

41. James MC, Bowman AS, Forbes KJ, Lewis F, McLeod JE, Gilbert L. Environmental determinants of Ixodes ricinus ticks and the incidence of *Borrelia burgdorferi* sensu lato, the agent of Lyme borreliosis, in Scotland. Parasitology. 2013 Feb;140(2):237–46.

42. Hanincová K, Schäfer SM, Etti S, Sewell HS, Taragelová V, Ziak D, et al. Association of *Borrelia afzelii* with rodents in Europe. Parasitology. 2003;126:11–20.

43. Milne A. Pasture Improvement and the control of the sheep tick *(Ixodes ricinus* L.). Ann Appl Biol. 1948;35:369–378.

44. Owen NW, Kent M, Dale P. Ecological effects of cultivation on the machair sand dune systems of the Outer Hebrides, Scotland. J Coast Conserv. 2000;6(2):155–70.

45. Gilbert L, Maffey GL, Ramsay SL, Hester A. The effect of deer management on the abundance of *Ixodes ricinus* in Scotland. Ecol Appl. 2012 Mar;22(2):658–67.

46. Hofmeester TR, Sprong H, Jansen PA, Prins HHT, Van Wieren SE. Deer presence rather than abundance determines the population density of the sheep tick, *Ixodes ricinus*, in Dutch forests. Parasites and Vectors. 2017;10(1):1–8.

47. Mysterud A, Easterday WR, Stigum VM, Aas AB, Meisingset EL, Viljugrein H. Contrasting emergence of Lyme disease across ecosystems. Nat Commun. 2016;7(6630):11882.

