## Appendix for "Emergence of Lyme disease on treeless islands in Scotland, UK"

**Corpus linguistic approach to extract common keywords from free text**

Corpus linguistic methods were used to explore the meanings in free-text responses of respondents to the question: “Do you think tick numbers and problems with ticks have changed over time?” The text responses were imported from MS Excel into notepad to form a corpus and then imported into a corpus linguistic analytical software tool (WordSmith Tools v7). Corpus linguistic analysis was performed as previously described (1).

The free-text responses were first analysed across all islands and then separated by high or low Lyme disease incidence island group. Prior to analysis, the words contained within each question were included in a “stop list” in the analysis for each question; this removed suggestion bias from the responses and prevents those words being identified as key words in the analysis.

Key word analysis identified key words in the grouped responses for each question. Key word analysis allows a useful set of preliminary words, each of which has a key-ness value assigned based on a comparison to the frequency of the word in the British National Corpus (BNC). The log-likelihood value was used as a measure of key-ness as per standard option in Wordsmith Tools (2). Key word clusters were also presented to present the meanings within each text.

Further exploration of the main key words using concordance and collocation analysis allowed further exploration of the context around the main key words. In other words, this contextual analysis revealed more of the meaning behind what respondents wrote.

**Habitat types of reported human tick bites**

Most participants (64.4%, n=333/517) provided information on the habitat type of their last tick bite and island of residence, with the most tick bites reported from heather moorland (39.3%, n=131/333), croft grassland (27.6%, n= 92/333), gardens (26.4%, n=88/333) and machair grassland (6.6%, n=22/333). There was a tendency for more tick bites in gardens and fewer in machair grassland in high LD incidence areas (gardens: Chi squared test, p = 0.053, 26/127 low LD incidence, 62/206 high LD incidence) (machair grassland: Chi squared test, p =0.036, 13/127 low LD incidence, 9/206 high LD incidence).

**Appendix References**

(1) Huntley SJ, Mahlberg M, Wiegand V, van Gennip Y, Yang H, Dean RS, et al. Analysing the opinions of UK veterinarians on practice-based research using corpus linguistic and mathematical methods. Prev Vet Med. 2018;150(January 2017):60–9.

(2) Dunning T. Accurate Methods for the Statistics of Surprise and Coincidence , Vol 19, No. 1, pp. 61-74. Comput Linguist. 1993;19(1):61–74

**Appendix Table 1.** Sampling site locations, *Ixodes ricinus* nymph density and *Borrelia burgdorferi* s.l. infection prevalence in study of environmental hazard in high and low Lyme disease (LD) incidence areas, Western Isles, UK.

| LD incidence | Island^*^ | Habitat^†‡^ | Year | Latitude/  Longitude | Total nymphs collected | Nymphs per 100m | Nymphs tested | Prevalence % |
| --- | --- | --- | --- | --- | --- | --- | --- | --- |
| Low | S. Harris | I. grassland | 2018 | 57.82297/-7.04121 | 100 | 32.5 | 100 | 0 |
|  |  |  | 2019 | 57.8033/-6.76844 | 190 | 20 | 50 | 2 |
|  |  |  | 2019 | 57.85990/-6.97867 | 1 | 0 | NA | NA |
|  |  |  | 2019 | 57.83909/-6.75505 | 6 | 0 | NA | NA |
|  |  | H. moorland | 2018 | 57.76642/-6.99558 | 100 | 10.5 | 100 | 0 |
|  |  |  | 2019 | 57.85464/-6.77910 | 9 | 1 | NA | NA |
|  |  |  | 2019 | 57.85119/-6.96178 | 21 | 1 | NA | NA |
|  |  |  | 2019 | 57.81542/-6.92514 | 1 | 0 | NA | NA |
|  | Barra | I. grassland | 2018 | 57.00991/-7.49054 | 98 | 5.5 | 98 | 0 |
|  |  |  | 2019 | 56.98296/-7.50262 | 1 | 0.5 | NA | NA |
|  |  |  | 2019 | 56.99873/-7.49991 | 1 | 0 | NA | NA |
|  |  |  | 2019 | 57.04221/-7.42730 | 1 | 0 | NA | NA |
|  |  | H. moorland | 2018 | 57.01508/-7.45006 | 110 | 6 | 100 | 0 |
|  |  |  | 2019 | 56.96077/-7.51683 | 59 | 4.5 | 54 | 0 |
|  |  |  | 2019 | 56.97012/-7.50559 | 137 | 8.5 | 57 | 0 |
|  |  |  | 2019 | 56.97535/-7.42723 | 84 | 2 | 50 | 6 |
| High | N. Uist | I. grassland | 2018 | 57.64579/-7.27850 | 18 | 1 | NA | NA |
|  |  |  | 2018 | 57.59542/-7.37803 | 59 | 44.5 | 50 | 18 |
|  |  |  | 2018 | 57.55027/-7.27865 | 51 | 17 | 50 | 2 |
|  |  |  | 2018 | 57.55679/-7.36161 | 52 | 1.5 | 49 | 6.12 |
|  |  | H. moorland | 2018 | 57.64992/-7.47042 | 52 | 17 | 50 | 4 |
|  |  |  | 2018 | 57.56901/-7.28658 | 15 | 0 | NA | NA |
|  |  |  | 2018 | 57.57699/-7.35361 | 57 | 48 | 50 | 14 |
|  |  |  | 2018 | 57.62612/-7.20569 | 76 | 4 | 76 | 5.26 |
|  | Benb. | I. grassland | 2018 | 57.41507/-7.30903 | 9 | 1 | NA | NA |
|  |  |  | 2018 | 57.42847/-7.35645 | 23 | 2.5 | NA | NA |
|  |  | H. moorland | 2018 | 57.43784/-7.36701 | 51 | 12 | 50 | 2 |
|  |  |  | 2018 | 57.46292/-7.29770 | 20 | 2 | 20 | NA |
|  | S. Uist | I. grassland | 2018 | 57.39698/-7.34315 | 9 | 0.5 | NA | NA |
|  |  |  | 2018 | 57.33157/-7.36658 | 0 | 0 | NA | NA |
|  |  |  | 2018 | 57.27389/-7.39276 | 76 | 10 | 58 | 12.07 |
|  |  |  | 2018 | 57.19942/-7.40313 | 76 | 10.5 | 50 | 4 |
|  |  |  | 2018 | 57.16089/-7.30559 | 251 | 58.5 | 77 | 9.09 |
|  |  |  | 2018 | 57.12438/-7.37993 | 35 | 2 | NA | NA |
|  |  | H. moorland | 2018 | 57.30218/-7.35176 | 53 | 34 | 50 | 6 |
|  |  |  | 2018 | 57.23865/-7.32935 | 50 | 31.5 | 50 | 0 |
|  |  |  | 2018 | 57.23901/-7.36996 | 55 | 6.5 | 50 | 10 |
|  |  |  | 2018 | 57.13368/-7.34022 | 50 | 7 | 50 | 2 |
|  |  |  | 2018 | 57.26320/-7.27952 | 50 | 36 | 50 | 2 |
|  |  |  | 2018 | 57.33028/-7.30772 | 76 | 69 | 76 | 5.26 |
|  |  |  | 2018 | 57.33718/-7.35609 | 25 | 2.5 | NA | NA |
|  |  |  | 2018 | 57.13750/-7.29402 | 9 | 0.5 | NA | NA |
|  | N. Uist | Bog,peatland | 2018 | 57.61533/-7.20634 | 51 | 1.5 | 50 | 2 |
|  |  |  | 2018 | 57.64040/-7.42523 | 56 | 4.5 | 50 | 14 |
|  |  |  | 2018 | 57.57971/-7.24579 | 21 | 1.5 | NA | NA |
|  |  |  | 2018 | 57.53646/-7.31195 | 50 | 15 | 50 | 0 |
|  |  | Garden | 2018 | sector 2 | 13 | 6.5 | NA | NA |
|  |  |  | 2018 | sector 3 | 11 | 5.5 | NA | NA |
|  |  |  | 2018 | sector 4 | 20 | 10 | NA | NA |
|  |  |  | 2018 | sector 5 | 2 | 1 | NA | NA |
|  |  | Machair | 2018 | 57.66823/-7.24728 | 0 | 0 | NA | NA |
|  |  |  | 2018 | 57.59891/-7.52762 | 0 | 0 | NA | NA |
|  |  |  | 2018 | 57.57246/-7.47268 | 0 | 0 | NA | NA |
|  | Benb. | Bog,peatland | 2018 | 57.46696/-7.33463 | 18 | 1 | NA | NA |
|  |  | Garden | 2018 | sector 7 | 11 | 5.5 | NA | NA |
|  |  | Machair | 2018 | 57.42549/-7.37725 | 0 | 0 | NA | NA |
|  | S. Uist | Bog,peatland | 2018 | 57.32618/-7.27926 | 50 | 4.5 | 50 | 0 |
|  |  |  | 2018 | 57.24386/-7.32181 | 59 | 10 | 50 | 18 |
|  |  |  | 2018 | 57.12952/-7.30258 | 36 | 2.5 | NA | NA |
|  |  |  | 2018 | 57.15575/-7.37453 | 50 | 2.5 | 50 | 0 |
|  |  |  | 2018 | 57.34627/-7.26833 | 76 | 8.5 | 50 | 0 |
|  |  |  | 2018 | 57.24486/-7.35349 | 270 | 15 | 50 | 10 |
|  |  |  | 2018 | 57.34174/-7.34557 | 32 | 1.5 | NA | NA |
|  |  |  | 2018 | 57.27817/-7.37005 | 4 | 0 | NA | NA |
|  |  | Garden | 2018 | sector 8 | 3 | 1.5 | NA | NA |
|  |  |  | 2018 | sector 9 | 64 | 12.5 | 50 | 6 |
|  |  |  | 2018 | sector 11 | 193 | 32.5 | 50 | 14 |
|  |  |  | 2018 | sector 12 | 100 | 17.5 | 56 | 1.79 |
|  |  |  | 2018 | sector 13 | 16 | 4 | NA | NA |
|  |  |  | 2018 | sector 14 | 73 | 36.5 | 49 | 16.33 |
|  |  |  | 2018 | sector 15 | 6 | 2.5 | NA | NA |
|  |  | Machair | 2018 | 57.35096/-7.39092 | 2 | 1 | NA | NA |
|  |  |  | 2018 | 57.30452/-7.39269 | 0 | 0 | NA | NA |
|  |  |  | 2018 | 57.24395/-7.42612 | 6 | 1 | NA | NA |
|  |  |  | 2018 | 57.15629/-7.40349 | 0 | 0 | NA | NA |

^*^Benbecula (Benb) North Uist (N. Uist), South Uist (S.Uist). South Harris (S. Harris).

^†^Improved grassland (I. grassland), Heather moorland (H. moorland).

^‡^Gardens were defined as areas next to a dwelling which were enclosed by a fence to restrict entry of livestock but not deer and ranged in size from 0.11 – 0.21 hectares. Gardens typically had a mowed lawn, with areas of shrubs, longer grass and trees. Latitude and longitude are not given for privacy reasons.

**Appendix Table 2.** Results of best fit generalized linear mixed model to explain variation in questing nymph density, infection prevalence and Lyme disease (LD) hazard among different habitat types sampled on high LD incidence islands. Differences in prevalence and LD hazard between different islands^*^ and habitats^†^ were tested.

| Model | Variable | Estimate | SE | p value | delta AIC^‡^ |
| --- | --- | --- | --- | --- | --- |
| Tick density | (Intercept) | 0.14 | 0.75 | <0.001 | NA |
|  | Habitat type  Heather moorland  Improved grassland  Bog and Peatland  Machair  Garden | (reference)  -0.34  -0.76  -3.32  0.17 | 0.51  0.49  0.80  0.51 | 0.511  0.121  <0.001  0.736 | 16.06 |
|  | Vegetation density | 0.14 | 0.04 | <0.001 | 13.68 |
|  | Humidity | -0.02 | 0.01 | 0.021 | 3.32 |
| Tick prevalence  (island) | (Intercept) | -3.01 | 0.25 | <0.001 | NA |
| Tick prevalence  (habitat) | (Intercept) | -3.00 | 0.28 | <0.001 | NA |
| LD hazard  (island) | (Intercept) | -5.52 | 0.42 | <0.001 | NA |
| LD hazard  (habitat) | (Intercept) | -5.13 | 0.43 | <0.001 | NA |

* Machair was excluded due to very low tick density at all sampled sites and garden sites were excluded from the model to test for differences between islands as gardens sampled on North Uist did not have the minimum sample size due to continuous dragging not being carried out at these sites. To test for differences among islands within high LD incidence areas, and habitat types excluding gardens and machair, 23 sites on North and South Uist, and grassland, moorland and bog and peatland sites were included.

^†^To test for differences in prevalence between gardens and other habitats, 18 sites on South Uist from garden, grassland, moorland and bog and peatland sites were included.

^‡^delta AIC is the change in AIC from removing each variable from the best fit model.

**Appendix Table 3.** Univariable analysis of factors affecting risk of tick bite exposure in residents of the Western Isles (classified as high, ≥5 tick bites a year, or low, < 5 tick bites a year)

| Variable | Responses | Sample size | Factor level | Odds ratio (95%CI) | p value^*^ |
| --- | --- | --- | --- | --- | --- |
| Age | 458 | 8/46  77/321  35/91 | 18-30 years  30-60 years  > 60 years | 1.00 (reference)  1.50 (0.70-1.60)  2.99 (1.30-7.52) | 0.009 |
| Gender | 460 | 85/318  37/142 | Female  Male | 1.00 (reference)  0.97 (0.61-1.51) | 0.88 |
| LD incidence | 455 | 34/196  87/259 | Low  High | 1.0 (reference)  2.41 (1.55-3.82) | <0.001 |
| Occupation risk | 437 | 77/331  15/48  27/58 | Indoor  Outdoor  Retired | 1.0 (reference)  1.50 (0.76-2.86)  2.87 (1.61-5.11) | 0.002 |
| Outdoor activity | 432 | 21/120 | < Most days  Most days | 1.0 (reference)  1.94 (1.16-3.37) | 0.011 |
| Cat/dog ownership | 460 | 37/129  85/331 | No cats or dogs  Cat and/or dog owner | 1.0 (reference)  0.86 (0.55-1.36) | 0.51 |
| Accessed information on ticks and Lyme disease | 453 | 7/34  114/419 | Not accessed  Accessed | 1.0 (reference)  1.44 (0.64-3.68) | 0.39 |
| Attitudes to risk from tick bites | 453 | 23/137  55/196  44/120 | Risks minor  Risks significant  Risks serious | 1.0 (reference)  1.93 (1.12-3.39)  2.87 (1.62-5.20) | 0.001 |
| Prevention measures against tick bites | 296 | 15/68  47/135  17/27  12/66 | None taken  Special clothing  Deer fence+/-other  Other | 1.0 (reference)  1.89 (0.98-3.80)  6.01(2.33-16.38  0.78 (0.33-1.83) | <0.001 |
| Frequency of checking for tick bites | 449 | 17/179  18/78  53/112  34/80 | <10%  11-50%  51-99%  100% | 1.0 (reference)  2.86 (1.38-5.95)  8.56 (4.68-16.34)  7.04 (3.66-14.01) | <0.001 |

*p value from likelihood ratio test compared to a null model.

**Appendix Table 4.** Results of best-fit general linear model to assess the effect of factors affecting the detection of any tick (live unfed, engorged, dead)^*^ within a home.

| Variable | Estimate | Standard error | p value | Odds ratio (95%CI) | delta AIC^†^ |
| --- | --- | --- | --- | --- | --- |
| Intercept | -0.72 | 0.24 | 0.003 | NA | NA |
| Cat or dog ownership  No cat or dog  Cat or dog owner | (reference)  1.4 | 0.23 | <0.001 | 4.07 (2.61-6.41) | 36.94 |
| Outdoor activity  < Most days  Most days | (reference)  0.51 | 0.23 | 0.028 | 1.67 (1.05-2.64) | 2.77 |

^*^ Tick presence in a home was reported commonly by 63.7% of respondents (274/424) who also gave information on other covariates in the full model. Live unfed ticks which pose a biting risk to humans was reported by 28.3% of these respondents (120/424).

^†^ delta AIC is the change in AIC from removing each variable from the best fit model.

**Appendix Table 5.** Results of general linear model to test for an association between whether tick numbers and associated problems are considered to be increasing over time in association with Lyme disease (LD) incidence (low, high).

| Variable | Estimate | Standard error | p value | Odds ratio (95%CI) | delta AIC^*^ |
| --- | --- | --- | --- | --- | --- |
| Intercept | 1.07 | 0.24 | <0.001 | NA | NA |
| LD incidence  Low  High | (reference)  1.5 | 0.4 | <0.001 | 4.46 (2.10-10.02) | 13.5 |

^*^ delta AIC is the change in AIC from removing each variable from the best fit model.

**Appendix Table 6.** Comparison of collocated words within high and low incidence locations for the top 3 keywords across all islands to the survey question ‘Do you think tick numbers and problems with ticks have changed over time?

| Keyword | Overall position  (Log Likelihood) | LD incidence | Collocated words | Collocated clusters (n) |
| --- | --- | --- | --- | --- |
| Deer | 1 (601) | High  Low | more, about, ticks,  numbers, garden, close  Sheep, ticks, more | More deer (7), Deer and (5), the deer (5), of deer (4), deer about (3), And Ticks (3), So deer (3)  The deer (4), deer are (3), on the (3) |
| Increased | 2 (445) | High  Low | Years, numbers, tick(s), last, significantly, definitely  Numbers, more, ticks, sheep | Have increased (12), definitely increased (5), increased significantly (4), increased over (4), increased dramatically (4), dramatically increased (4), moorland increased (3)  Have increased (14), to have (5), they have (4), seem to (3) |
| Sheep | 3 (272) | High  Low | Dipping, dipped, no  Deer, dipping, numbers, increased | Sheep, dipping (3)  Sheep dipping (3), the sheep (3), on the (3) |

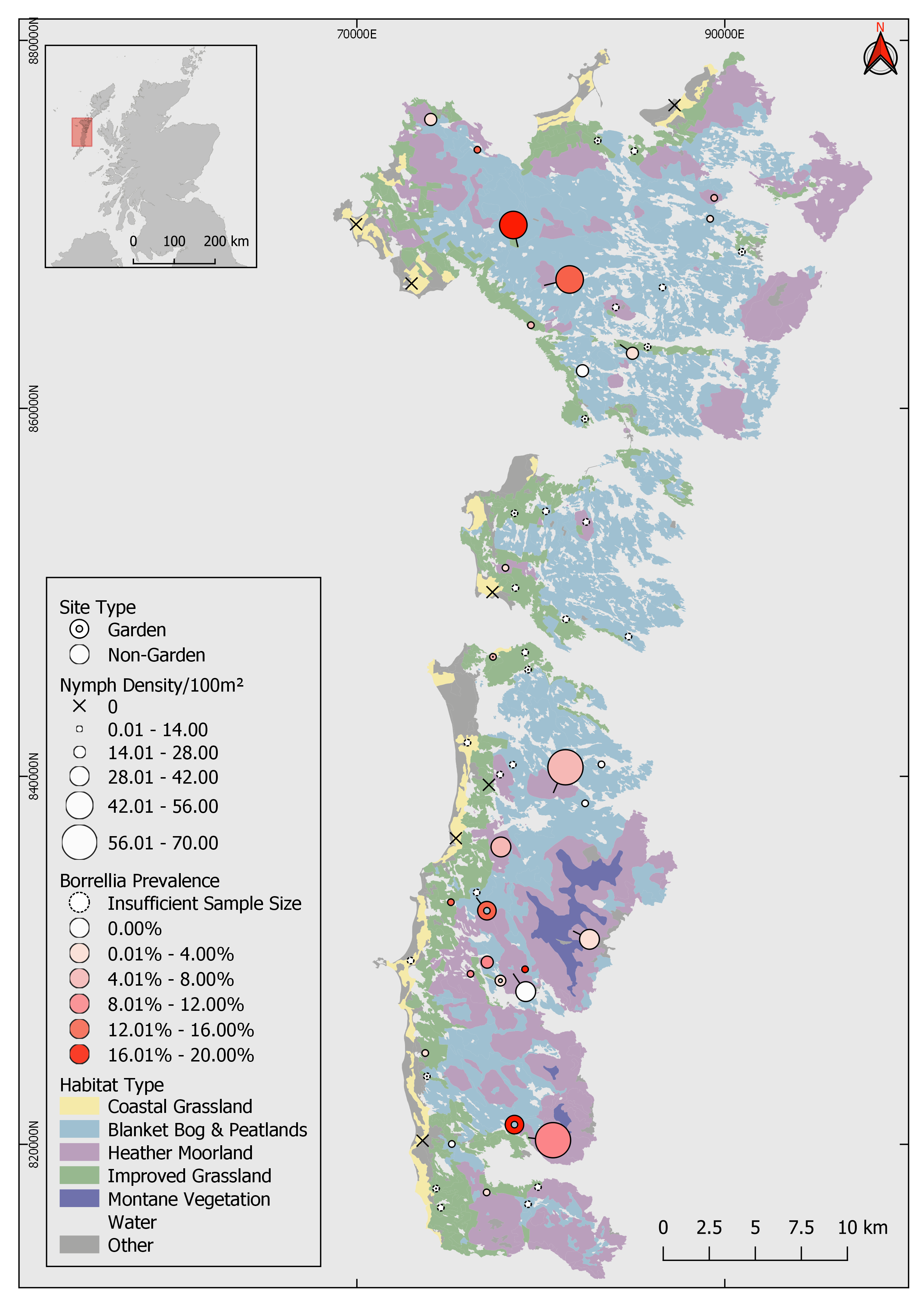

**Appendix Figure 1.** Map of study sites in improved grassland, heather moorland, bog and peatland, machair grassland and domestic gardens within high Lyme disease incidence islands in the Western Isles, UK sampled in 2018. Circles are in proportion to the questing tick density and *Borrelia burgdorferi* sensu lato prevalence is represented by the graded colour of the circles. Prevalence was not estimated at sites where < 50 ticks were collected. Sites at which no ticks were collected from are shown as a ‘x’. Major habitats which were sampled are shown on the map and sectors for stratified sampling are shown as a grid with numbers alongside the map.

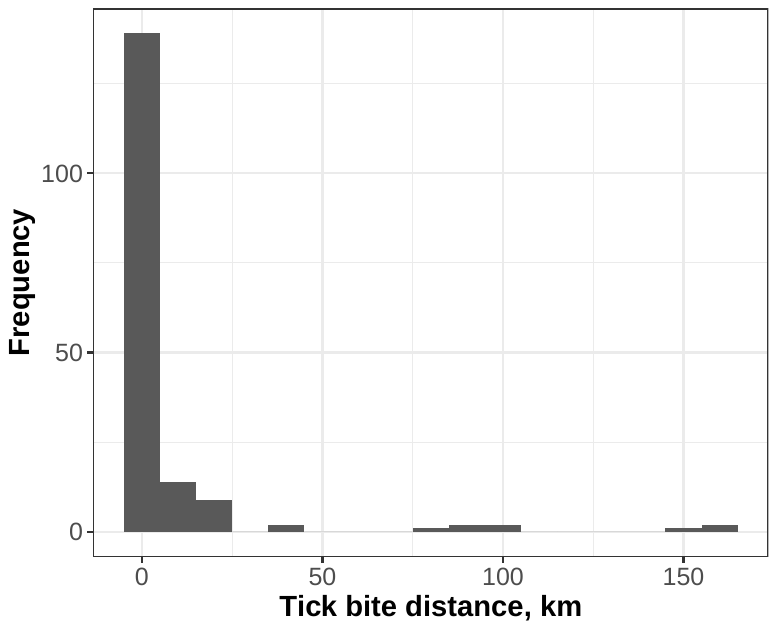

**Appendix Figure 2.** Plot of distribution of tick bite distance from the home address, from survey of residents in the Western Isles, UK, 2018.
